# Prefrontal cortical connectivity mediates locus coeruleus noradrenergic regulation of inhibitory control in older adults

**DOI:** 10.1101/2021.06.29.450427

**Authors:** Alessandro Tomassini, Frank H. Hezemans, Rong Ye, Cam-CAN, Kamen A. Tsvetanov, Noham Wolpe, James B. Rowe

**Affiliations:** MRC Cognition and Brain Sciences Unit, University of Cambridge, UK; Department of Clinical Neuroscience and Cambridge University Hospitals NHS Trust, University of Cambridge, UK; Centre for Speech, Language and the Brain, Department of Psychology, University of Cambridge, UK

**Keywords:** Locus Coeruleus, neuromelanin, response inhibition, stop-signal task, healthy ageing, functional connectivity

## Abstract

Response inhibition is a core executive function enabling adaptive behaviour in dynamic environments. Human and animal models indicate that inhibitory control and control networks are modulated by noradrenaline, arising from the locus coeruleus. The integrity (i.e., cellular density) of the locus coeruleus noradrenergic system can be estimated from magnetization transfer sensitive magnetic resonance imaging, in view of neuromelanin present in noradrenergic neurons of older adults. Noradrenergic psychopharmacological studies indicate noradrenergic modulation of prefrontal and frontostriatal stopping-circuits in association with behavioural change. Here we test the noradrenergic hypothesis of inhibitory control, in healthy adults. We predicted that locus coeruleus integrity is associated with age-adjusted variance in response inhibition, mediated by changes in connectivity between frontal inhibitory control regions. In a preregistered analysis, we used magnetization transfer MRI images from N=63 healthy adults aged above 50 years who performed a stop-signal task, with atlas-based measurement of locus coeruleus contrast. We confirm that better response inhibition is correlated with locus coeruleus integrity and stronger connectivity between pre-supplementary motor area and right inferior frontal gyrus, but not volumes of the cortical regions. We confirmed a significant role of prefrontal connectivity in mediating the effect of individual differences in the locus coeruleus on behaviour, whereby this effect was moderated by age, over and above adjustment for the mean effects of age. Our results support the hypothesis that in normal populations, as in clinical settings, the locus coeruleus noradrenergic system regulates inhibitory control.

## Introduction

Response inhibition underpins the control of everyday behavior, and is impaired in many neurological and psychiatric disorders (Passamonti, Lansdall, & Rowe, 2018). There is converging evidence from animal models (Bari et al., 2011; Bari & Robbins, 2013; Eagle & Baunez, 2010) and human psychopharmacology (Chamberlain et al., 2006; Chamberlain et al., 2009; Robinson et al., 2008), that the noradrenergic system facilitates inhibitory control for action cancellation. A prefrontal cortical network is also implicated in such response inhibition, including the inferior frontal gyrus (rIFG) and pre-supplementary motor area (preSMA; Aron, Robbins, & Poldrack, 2014; Chambers et al., 2006; Duann, Ide, Luo, & Li, 2009; Forstmann et al., 2012; Frank, Scheres, & Sherman, 2007; Rae, Hughes, Anderson, & Rowe, 2015).

Cerebral noradrenergic innervation arises from the locus coeruleus, in the brainstem. By exploiting the accumulation of iron-rich neuromelanin, magnetization-transfer (MT) sensitive magnetic resonance imaging sequences can be used to assess the integrity of the locus coeruleus (Liu et al 2019; Ye et al., 2020; but see also Watanabe et al., 2019). Such studies have associated locus coeruleus integrity with diverse cognitive function in health (Liu et al., 2020) and disease (Dahl et al., 2020; Holland, Robbins, & Rowe, 2021; O’Callaghan et al., 2021; Ye et al., 2021). Phasic and tonic activities in the locus coeruleus have been proposed to afford behavioural flexibility and inhibitory control (Aston-Jones & Cohen, 2005; Bouret & Sara, 2005; Dayan & Yu, 2006; Passamonti, Lansdall, & Rowe, 2018). This might be achieved by modulation of connectivity within the prefrontal network (Chambers et al., 2006; Duann et al., 2009; Forstmann et al., 2012; Aron et al., 2014; Rae et al., 2015; Ye et al., 2015; Rae et al., 2016). Here, we tested this hypothesis by linking locus coeruleus integrity to functional connectivity between rIFG and preSMA, as a predictor of behaviour.

Previous analysis of cortical connectivity in healthy adults showed that the influence of cortical connectivity on inhibitory control differs with age, such that efficient performance in older adults relies more strongly on connectivity than in their younger counterparts (Tsvetanov et al., 2018).

Moreover, sub-regions of the locus coeruleus have different projection distributions (Mason & Fibiger, 1979; Loughlin, Foote, Bloom, 1986) with differential associations to cognition, behaviour and pathology. For example, *in vivo* evidence suggest greater degeneration of the caudal sub-region of the locus coeruleus compared to central and rostral sub-regions in Parkinson’s disease (O’Callaghan et al., 2021). Healthy ageing and age-related cognitive decline are more strongly associated with changes in the rostral locus coeruleus (Betts, Cardenas-Blanco, Kanowski, Jessen, & Düzel, 2017; Liu et al., 2020; Dahl et al., 2020).

We hypothesized that variations in the locus coeruleus integrity would drive noradrenergic-dependent changes in inhibitory control, over and above the main effect of age on the locus coeruleus. We tested whether such relationship would vary across locus-coeruleus sub-regions and across different ages. In this pre-registered cross-sectional study, we used a 3-D MT weighted MRI sequence to assess the relationship between locus coeruleus integrity *in vivo* and test its relationship with inhibitory control in cognitively normal healthy adults from the Cambridge Centre for Ageing and Neuroscience cohort (Cam-CAN; Shafto et al., 2014) using the Stop-Signal Task (SST). Our study has four main advances with respect to an earlier analysis of the Cam-CAN cohort (Liu et al., 2020). First, we use an atlas-based segmentation of the locus coeruleus, which provides unbiased estimation, with good accuracy and reliability compared to manual and semi-automatic segmentation approaches (Ye et al., 2021). Second, we focus on middle-aged and older healthy adults, because neuromelanin accumulates with age (Zecca, Youdim, Riederer, Connor, & Crichton, 2004) and in younger participants the locus coeruleus neurons may not yet be sufficiently pigmented to allow reliable inference on structural integrity by MT weighted MRI. Third, we follow the new consensus recommendations for estimating the stop-signal reaction time (SSRT, Verbruggen et al.,2019), and use hierarchical Bayesian estimation of a parametric ex-Gaussian race model of the stop-signal task which enables isolating attentional confounds from the estimation of SSRTs (Matzke, Dolan, Logan, Brown, & Wagenmakers, 2013). Fourth, we examine whether locus coeruleus integrity is related to modulation of connectivity within the prefrontal stopping-network quantified by psychophysical interactions measures that reflect response inhibition-related changes in the connectivity between different areas (Friston et al., 1997; Tsvetanov et al., 2018).

## Materials and Methods

### Preregistration

Before data analysis, we preregistered our analyses, sample size, variables of interest, hypotheses, procedures for data quality checking and data analysis procedures in the Open Science Framework. The preregistered information, code and data to reproduce manuscript figures are available through the Open Science Framework (<https://osf.io/zgj9n/>).

### Participants

We used data from the “Stage 3” cohort in the Cambridge Centre for Aging and Neuroscience population-based study of the healthy adult life span (See Shafto et al, 2014 for details). Within this cohort, we focused on 114 participants (18-88years) who performed a stop-signal task during fMRI. None of the Cam-CAN participants had a diagnosis of dementia or mild cognitive impairment, and all scored above consensus thresholds for normal cognition on the ACE-R (>88/100). After quality control of behavioural data and MRI scans, there were 63 datasets from participants aged 50 years or older (see table 1 for details). Note that our primary statistical inferences are Bayesian, where inferences are based on relative evidence for alternate models, rather than testing a null hypothesis alone. However, for secondary frequentist statistics where type I or type II error may arise, we computed the achieved power using G*Power 3.1. With a nominal alpha level=0.05, N=63 provides 86% power to test the interaction (moderation) term of a multiple linear regression with five predictors, assuming a medium-sized effect (*f*^*2*^=0.15).

**Table 1.** Demographics

| Variables | Overall |
| --- | --- |
| N | 63 |
| Age (median [range]) | 68.00 [50.00, 88.00] |
| Sex | 33M/30F |
| ACE-R (median [range]) | 95.00 [89.00, 100.00] |

### Procedure

The stop-signal task assesses cognitive control systems involved in action cancellation using stop-signal trials (n =80; approximately 50% of which were successful), randomly interleaved among go trials (n = 360) and no-go trials (n = 40) during two consecutive scanning runs (Figure 1A). On go trials participants saw a black arrow (duration 1000ms) and indicated its direction by pressing left or right buttons with the index or middle finger of their right hand. On stop-signal trials, the black arrow changed color (from black to red) concurrent with a tone, after a short, variable “stop-signal” delay (SSD). On no-go trials, the arrow was red from the outset (i.e., SSD = 0), along with a concurrent equivalent tone. Participants were instructed to withhold button pressing if the arrow was red or became red. The length of the SSD varied between stop-signal trials in steps of 50ms, and was titrated to participants’ performance using an on-line tracking algorithm to target a 50% successful response cancellation. No-go trials were included as stop trials with nominal SSD = 5ms (Matzke, Curley, Gong & Heathcote, 2019).

**Figure 1.**
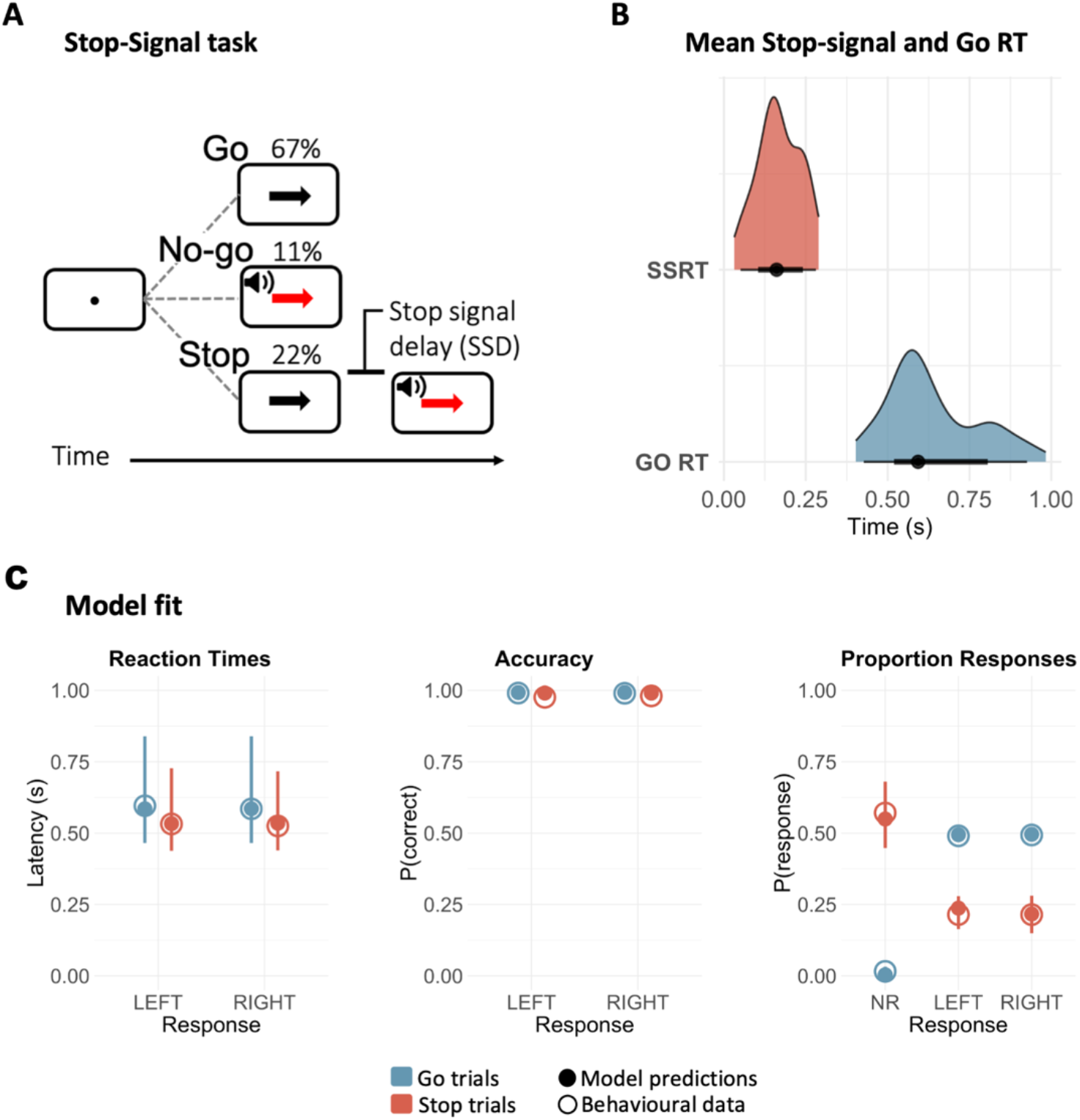
Stop-Signal task (A) and SSRTs estimated by the ex-Gaussian race model of response inhibition (B). A) In the Stop-Signal task, participants respond to the direction of a black arrow by pressing the corresponding key as accurately and as quickly as possible. Occasionally, a red arrow and a tone (stop signal) require the participants to inhibit their response. The stop signal could either appear immediately after the fixation point (no-go trials), or after a short delay (stop signal delay) that varies across trials. B) Distributions of mean SSRT and go RT. The ex-Gaussian race model depicts task performance as a race between a stop process and a go process. Successful inhibition in stop and no-go trials occurs when the stop process finishes its race before the go process. The black circles indicate the medians, the thick black segments depict the 66% quantile intervals, and the thin black segments depict the 95% quantile interval. C) Posterior predictive checks: Comparing empirical data (open circles) to simulated results from the fitted model (filled circles). Within each panel, the group-level median values are plotted separately for each response (left, right and NR -no response – when applicable) and trial type (go, stop). Please note that for Reaction Times and Accuracy, responses in stop trials correspond to commission errors, whereas for Proportion Responses, no response in go trials are omission errors. Model predictions are represented by the median (filled circles) and 95% quantile intervals (error bars) of 100 simulated participants, randomly drawn from the joint posterior distribution.

### Imaging

Imaging data were acquired with a 3T Siemens TIM trio with a 32-channel head coil. For each participant, a 3D structural MRI was acquired using a T1-weighted sequence with generalized autocalibrating partially parallel acquisition. The adopted parameters were as follows: acceleration factor, 2; repetition time (TR) = 2250 ms; echo time (TE) = 2.99 ms; inversion time = 900 ms; flip angle = 9°; field of view (FOV) = 256 × 240 × 192 mm; resolution = 1 mm isotropic; acquisition time = 4 min 32 s. For fMRI, echoplanar imaging (EPI) captured 32 slices in sequential descending order with slice thickness of 3.7 mm and a slice gap of 20% for whole-brain coverage. The adopted parameters were as follows: TR = 2000 ms; TE = 30 ms; flip angle = 78°; FOV = 192 × 192 mm; resolution = 3 × 3 × 4.4 mm, with a total duration of ∼ 10 min 30 s. For preprocessing details, see Taylor et al. (2017).

Image processing followed a co-registration pipeline similar to Ye et al., 2020, with Advanced Normalization Tools (ANTs v2.2.0) software and in-house Matlab scripts. MT images were N4 bias field corrected for spatial inhomogeneity (number of iterations at each resolution level: 50×50×30×20, convergence threshold: 1×10-6, isotropic sizing for b-spline fitting: 200) and to skull-strip T1-w images after segmentation and reconstruction (SPM12 v7219, www.fil.ion.ucl.ac.uk/spm/software/spm12). The resulting T1-w and pre-processed MT-weighted images were entered into a T1-driven, crossmodality coregistration pipeline to warp the individual MT and MT-off images to the isotropic 0.5 mm ICBM152 (International Consortium for Brain Mapping) T1-w asymmetric template.

We created an unbiased study-specific T1-w structural template using individual skull-stripped T1-w images from all participants (Figure 2A). Native T1-w images were first rigid and affine transformed, and then processed with a hierarchical nonlinear diffeomorphic step at five levels of resolution, repeated by six runs to improve convergence. The resulting T1-w group template was then registered to the standard ICBM152 T1-w brain. Four steps of deformations were estimated in the following order: MT-off to MT, T1-w to MT-off, T1-w to T1-w group template and T1-w group template to ICBM152 T1-w template. The resulting parameters were used as the roadmap for MT image standardization to the ICBM brain in one step. For co-registration details, see Ye et al., (2020).

**Figure 2.**
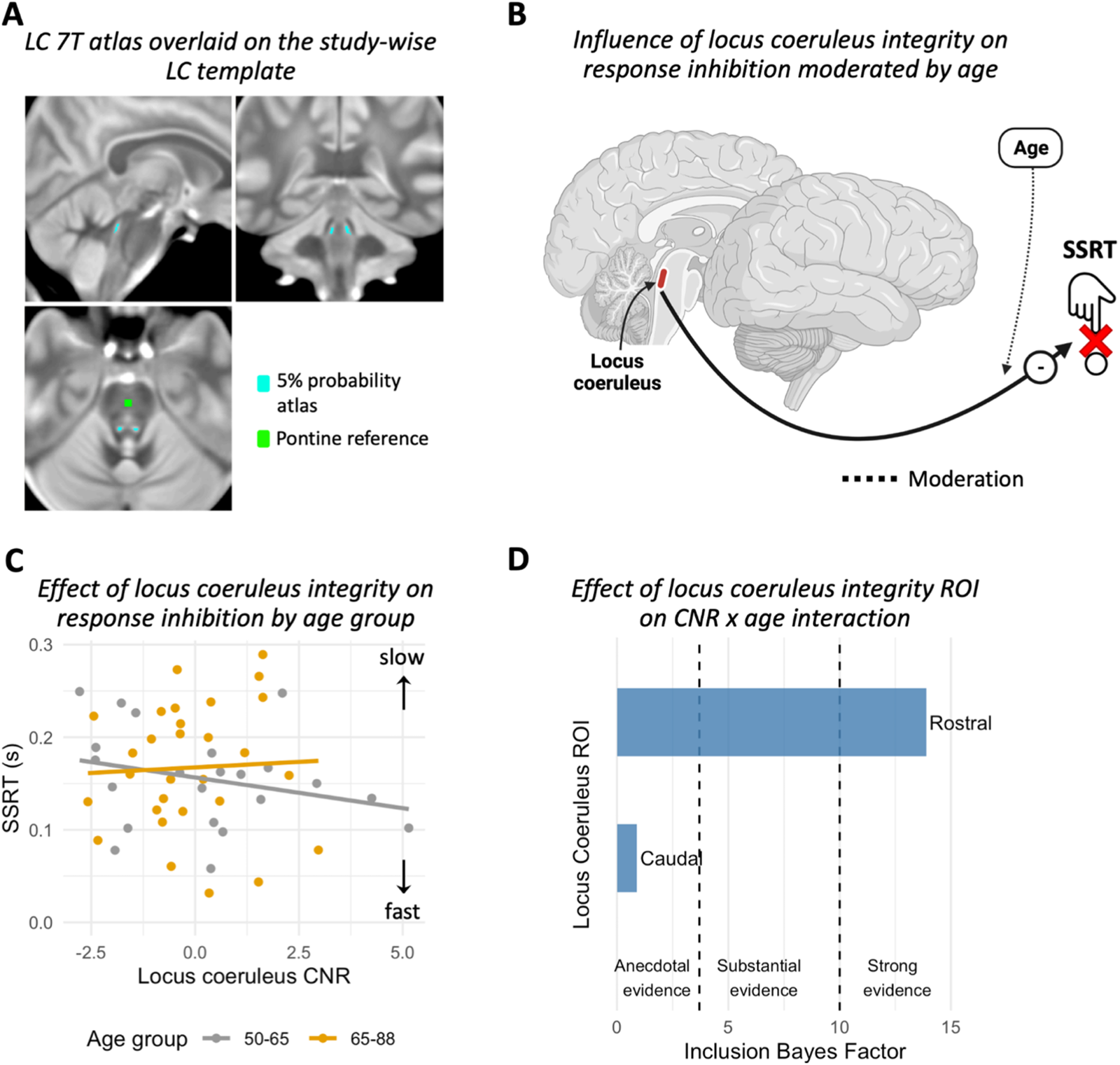
A) Study specific atlas of the locus coeruleus (light blue) and reference region in the central pons (green). B-C) SSRT estimates as a function of age adjusted locus coeruleus contrast-to-noise ratio (CNR) and age group. The interaction with age is estimated as a continuous variable (see text) but binarized for visualization purposes only. D) Bayesian evidence for an association between the integrity of rostral, and caudal sub-regions of the locus coeruleus with response inhibition.

To facilitate accurate extraction of the locus coeruleus signal we adopted a probabilistic locus coeruleus atlas (Ye et al., 2020) generated from ultra-high resolution 7T data accompanied by a multi-modality co-registration pipeline. As a measure of locus coeruleus integrity, we quantified contrast by calculating the contrast-to-noise ratio (CNR) with respect to a reference region in the central pons (Figure 2A). A CNR map was computed voxel-by-voxel on the average MT image for each subject using the signal difference between a given voxel (*v*) and the mean intensity in the reference region (Mean_REF_) divided by the standard deviation (SD_REF_) of the reference signals

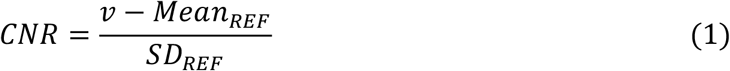

CNR values were computed bilaterally on the CNR map by applying the independent locus coeruleus probabilistic atlas (5% probability version to improve sensitivity). To obtain summary indices of CNR for subsequent analyses, we extracted mean CNR values for the rostral, middle and caudal portions of the LC, collapsing across the left and right locus coeruleus.

Voxel-based morphometry estimates of grey matter volume for rIFG and preSMA cortical areas were extracted from functionally-defined masks previously defined in Tsvetanov et al., 2018. For the fMRI stop-signal task-related functional connectivity we used psychophysiological interaction measures between rIFG and preSMA as estimated in Tsvetanov et al., 2018. Specifically, group independent components analysis (ICA) decomposed the fMRI signal into functional components that were activated in the contrast *successful stop > unsuccessful stop*. Then correlational psychophysiological interactions analysis was used to estimate patterns of functional connectivity modulation between rIFG and preSMA. The resulting measure quantifies differences in connectivity (i.e., modulation) between successful and unsuccessful action cancellation trials within the rIFG-preSMA network (see Tsvetanov et al., 2018 for further details).

### Analyses

To infer the latency of the unobservable response inhibition (i.e., the stop-signal reaction time; SSRT) we adopted hierarchical Bayesian estimation of a parametric race model of the stop-signal task (Matzke, Dolan, Logan, Brown, & Wagenmakers, 2013). Accordingly, performance on the stop-signal task is modeled as a race between three independent processes: one corresponding to the stop process, and two corresponding to go processes that match or mismatch the go stimulus. Successful response inhibition in stop-signal trials occurs when the stop process finishes its race before both go processes. Correct responses on go trials, instead, require the matching go process to finish its race before the mismatching go process. The finish time distribution of the stop process is inferred by estimating the RT distribution of unsuccessful stop trials (i.e., signal respond RTs; see Matzke, Dolan, Logan, Brown, and Wagenmakers, 2013 for details). The model assumes that the finish times of the stop and go processes follow an ex-Gaussian distribution (Heathcote et al., 2018). Thus, for each process, we described the corresponding ex-Gaussian distribution by estimating the mean μ and standard deviation σ of its Gaussian component, and the mean (i.e., inverse rate) τ of its exponential component.

Further, we estimated the probability that the stop and go processes failed to start, referred to as “trigger failure” and “go failure” (Matzke, Curley, Gong, & Heathcote, 2019). These attentional failures can be common in the stop-signal task and, if not accounted for, bias estimation of the stop process (Band et al., 2003; Matzke et al., 2019; Skippen et al., 2019). Prior to fitting the model, we excluded implausibly fast (< 0.25s) RTs, as well as outliers go RTs exceeding ±2.5 standard deviations from the participant’s mean (Matzke, Dolan, Logan, Brown, & Wagenmakers, 2013; O’Callaghan et al., 2021).

We estimated the posterior distributions of the parameters using Markov Chain Monte Carlo (MCMC) sampling. The parameters were estimated hierarchically, such that parameters for a given participant are assumed to be drawn from corresponding group-level normal distributions. We adopted prior distributions identical to those suggested by the model developers (Heathcote et al., 2018), except for slightly higher prior mean values for μ_go-match_ (1.5s), μ_go-mismatch_ (1.5s) and μstop (1s), to account for slower RT in older age (O’Callaghan et al., 2021). MCMC sampling initially ran with 33 chains (i.e., three times the number of parameters), with thinning of every 10th sample and a 5% probability of migration. Visual inspection of the MCMC chains as well as the potential scale reduction statistic 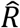 (less than 1.1 for all parameters; mean 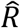 across subjects mean±sd: 1.009±0.002) were used to assess model fit convergence. After confirming convergence, an additional 500 iterations for each chain were run to obtain a posterior distribution for each parameter. The model’s goodness of fit was confirmed by comparing the observed data to simulated data generated from the model’s posterior predictive distribution (Figure 1C).

The SSRT was the primary outcome of interest, computed as the mean of the ex-Gaussian finish time distribution of the stop process, which is given by μ_stop_ + τ_stop_. We repeated this computation for each MCMC sample to approximate a posterior distribution of SSRT. The same approach was adopted to draw a posterior distribution of go RT (μ_go-match_ + τ_go-match_).

We used the statistical software package R (http://www.r-project.org/), following our preregistered analysis plan (<https://osf.io/zgj9n/>). For primary analyses we used Bayesian statistics which enables to quantify evidence in favor of the alternate hypothesis as well as evidence for the null hypothesis (of an absence of effect). We quantify relative evidence through Bayes Factors (BF) and for their interpretation we adhere to consensus guidelines (Jeffreys, 1961). For completeness, we also present classical frequentist analyses with α = 0.05 criterion for significance.

For all the regression analyses, we included sex as a binary covariate to account for possible sex-related differences in locus coeruleus signal (Clewett et al., 2016). We additionally controlled for changes in grey matter volume of the rIFG and preSMA, and check that regression assumptions are met. We include age as a continuous moderator to take into account possible age-related changes in the reliance of response inhibition on both locus coeruleus integrity and connectivity (Tsvetanov et al., 2018). To prevent collinearity issues, the continuous variables forming the interaction term of regression were mean centered. Moreover, since we are focusing on changes over and above the main effect of age on the locus coeruleus, locus coeruleus CNR values were regressed onto age and the residuals used as predictor.

For the moderated mediation analysis, the choice of the model structure was guided by a previous interventional study of the effect of atomoxetine on functional connectivity between preSMA and rIFG (Rae et al., 2016, see Figure 4a for a depiction of the model). The approach allows us to estimate the shared variance between locus-coeruleus signal and functional connectivity. The model was tested using the PROCESS macro model 15 in R using a bootstrap approach (Preacher and Hayes, 2004). The moderated mediation model tests for locus coeruleus-induced variability in response inhibition (direct effect, path *c*), and in functional connectivity mediating the effect on response inhibition (mediated effect, path *ab*). The model also tests whether the direct and indirect (i.e., mediated) effects of locus coeruleus on response inhibition change with age (age moderation, paths *b2* and *c2*).

### Software and Equipment

The ex-Gaussian model fitting was performed with the Dynamic Models of Choice toolbox (Heathcote et al., 2019), implemented in R (version 4.0, R Core Team, 2019). Further statistical analyses in R used the ‘tidyverse’ (Wickham et al., 2019) for data organization and visualization, ‘processR’ with ‘lavaan’ (Rosseel, 2012) packages for path analysis, and the ‘BayesFactor’ (Morey et al., 2018) and ‘bayestestR’ (Makowski, Ben-Shachar, and Lüdecke, 2019) packages for Bayes Factor analysis. Figures 2B, 3A, 4A were created with BioRender.com.

## Results

We confirmed the task performance expectations in that (i) the group SSRT (median 161ms) and GoRT (median 593ms) were within the range expected from the literature on related tasks, as show in in figure 1B; (ii) Commissions error latencies were shorter than accurate GoRT, indicative of impulsive responses, figure 1C; and (iii) errors and latencies were approximately equal between left and right hand responses, figure 1C. The Stop signal algorithm converged on average 57.4% accuracy (SD 12.8%). In the following sections, we relate individual differences in performance to locus coeruleus CNR, using the 5% atlas, but note that results are qualitatively similar using the more conservative atlas threshold of 25%.

### Locus coeruleus integrity predicts individual differences in response inhibition

We tested for effects of locus coeruleus integrity (as CNR) on response inhibition (the SSRT) using multiple linear regression (Figure 2B). The locus coeruleus integrity was associated with faster inhibitory responses but the strength of such association diminishes with age (Figure 2C; interaction LC CNR × age: BF_H1_ = 16.866; t_55_ = 2.993; p = 0.004). This moderation remains significant even after controlling for loss of grey matter in preSMA and rIFG regions, crucial components of the stopping-network (BF_H1_ = 19.068; t_55_ = 3.08; p = 0.003). No significant main effects were observed for sex (BF_H1_ = 0.407; t_55_ = −0.767; p = 0.446) nor for volumetric differences in the preSMA (BF_H1_ = 0.714; t_55_ = - 0.817; p = 0.417) and rIFG (BF_H1_ = 0.623; t_55_ = 0.366; p = 0.715) across subjects. To confirm that the parameter estimates of the multiple linear regression were not driven by the excessive influence of any given participant, we calculated Cook’s distance which quantifies the influence of each data point on all parameters of the linear model simultaneously. As a rule of thumb, a participant is deemed overly influential if their Cook’s distance exceeds 4 divided by the total number of participants (Van der Meer, Te Grotenhuis, Pelzer, 2010). In our sample, the Cook’s distance values ranged from 0 to 0.056, within the applicable cut-off value of 4/62 = 0.64. In a further effort to ensure robustness of our results, we re-fit the model using robust linear regression, which is less sensitive to data outliers than the conventional ordinary least squares method. The results of the robust analysis confirmed the results from the conventional linear regression showing a significant locus coeruleus integrity × age interaction (t_55_ = 3.15; p = 0.0025), but no significant main effects of sex (t_55_ = - 0.618; p = 0.538) or grey volume in preSMA (t_55_ = - 0.811; p = 0.414) and rIFG (t_55_ = - 0.501; p = 0.616).

The observed moderating effect of age on the association of locus coeruleus integrity with inhibitory control might be confounded by differences in the degree to which a participant has “successfully aged” cognitively and maintained cognitive ability on par with early adult life in contrast to “unsuccessfully aged” cognitively with decline in cognitive ability. To estimate this change in cognitive ability, one can compare current fluid intelligence to crystallized intelligence. This difference (ability discrepancy score) approximates the degree to which a participant has sustained or changed their cognitive ability, with lower fluid than crystallized intelligence as a marker of “unsuccessful aging” (McDonough et al., 2016). We tested for a possible role of unsuccessful ageing by including ability discrepancy in the interaction term locus coeruleus CNR × age, while controlling for sex and cortical volumetric changes. Bayesian analysis confirmed the age-moderated relationship between locus coeruleus CNR and response inhibition observed in the previous analyses (BF_H1_ = 12.580; although not significant with frequentist analysis t_51_ = 1.567; p = 0.123). However, there was no significant main effect of ability discrepancy (BF_H1_ = 0.446; t_51_ = −0.482; p = 0.632), nor interaction effects of ability discrepancy × age (BF_H1_ = 0.497; t_51_ = 0.071; p = 0.944), CNR × ability discrepancy (BF_H1_ = 0.775; t_51_ = 1.053; p = 0.297), or CNR × ability discrepancy × age (BF_H1_ = 1.753; albeit marginally significant with frequentist analysis t_51_ = 2.017; p = 0.049). Therefore, age affects the modulatory effect of locus coeruleus integrity on response inhibition irrespective of individuals’ lifetime decline in cognitive ability.

To probe the specificity of the relationship between locus coeruleus integrity and action cancellation, we repeated the linear regression analysis replacing SSRTs with successful *Go* reaction times as outcome. There was evidence for slowing of Go reaction times with age (BF_H1_ = 6.824; t_55_ = 2.714; p = 0.0088) but no evidence for a link between Go reactions times and locus coeruleus integrity (main effect CNR: BF_H1_ = 0.366; t_55_ = 0.602; p = 0.549; interaction CNR × age: BF_H1_ = 0.617; t_55_ = 0.904; p = 0.369).

### Integrity of the rostral sub-region of the locus coeruleus is associated with response inhibition

Sub-regions of the locus coeruleus may have differential associations to cognition, behaviour and pathology given the heterogeneity in its topographic organization. Here we estimated the association between the integrity of sub-regions of the locus coeruleus with response inhibition. In the pre-registered analysis plan, we proposed to fit a multiple linear regression model including the interaction between rostral, middle, caudal subregions of the locus coeruleus and age while controlling for sex and gray matter volume as in our previous analyses. However, diagnostic tests indicated a strong multicollinearity in the interaction term driven by the middle CNR (variance inflation factor > 10). We therefore deviate from the pre-registered plan for this test, to exclude the middle sub-region from the model. The multicollinearity issue was solved (variance inflation factor < 10) and all the linear regression assumptions (e.g., skewness, kurtosis, heteroscedasticity) were met.

The multiple linear regression results show strong evidence for an association between response inhibition and the rostral sub-region mediated by age (rostral CNR × age: BF_H1_ = 13.905; t_50_ = 2.735; p = 0.0086) (Figure 2D). There were no significant interaction effects for CNR × age (BF_H1_ = 0.855; t_50_ = −1.090; p = 0.280), caudal CNR × rostral CNR (BF_H1_ = 0.607; t_50_ = 0.715; p = 0.477) and caudal CNR × rostral CNR × age (BF_H1_ = 0.545; t_50_ = −0.272; p = 0.786) nor for the main effects of caudal CNR (BF_H1_ = 0.658; t_50_ = −1.553; p = 0.126), sex (BF_H1_ = 0.634; t_50_ = −1.281; p = 0.206) and grey matter volume in preSMA (BF_H1_ = 1.345; t_50_ = −1.463; p = 0.149) and rIFG(BF_H1_ = 0.824; t_50_ = −0.350; p = 0.728).

### Modulation of connectivity between preSMA and rIFG predicts individual differences in response inhibition

We tested whether the performance related connectivity (between successful and unsuccessful stop trials) within the prefrontal stopping-network was related to individual differences in response inhibition (SSRT), using multiple linear regression. Age moderated the relationship between preSMA-rIFG connectivity and response inhibition (BF_H1_ = 4.644; t_55_ = 2.311; p = 0.024) (Figure 3B) qualitatively similar to the effect observed when locus coeruleus CNR was used as predictor. There were no significant main effects for sex (BF_H1_ = 0.449; t_55_ = −0.948; p = 0.347) or volumetric grey matter differences in preSMA (BF_H1_ = 0.566; t_55_ = −0.494; p = 0.623) and rIFG regions (BF_H1_ = 0.533; t_55_ = −0.238; p = 0.812). We confirmed robustness of our linear regression results by examining the Cook’s distance of each participant (range 0 – 0.058, below the 0.64 threshold) as well as by showing that results were qualitatively identical when re-fitting the data with a robust regression approach. Specifically, the robust regression results confirm a significant Connectivity × age interaction (t_55_ = 2.754; p = 0.008) but no significant main effects for sex (t_55_ = −0.622; p = 0.536) or grey matter volume in in preSMA (t_55_ = −0.507; p = 0.536) and rIFG (t_55_ = −0.349; p = 0.724).

**Figure 3.**
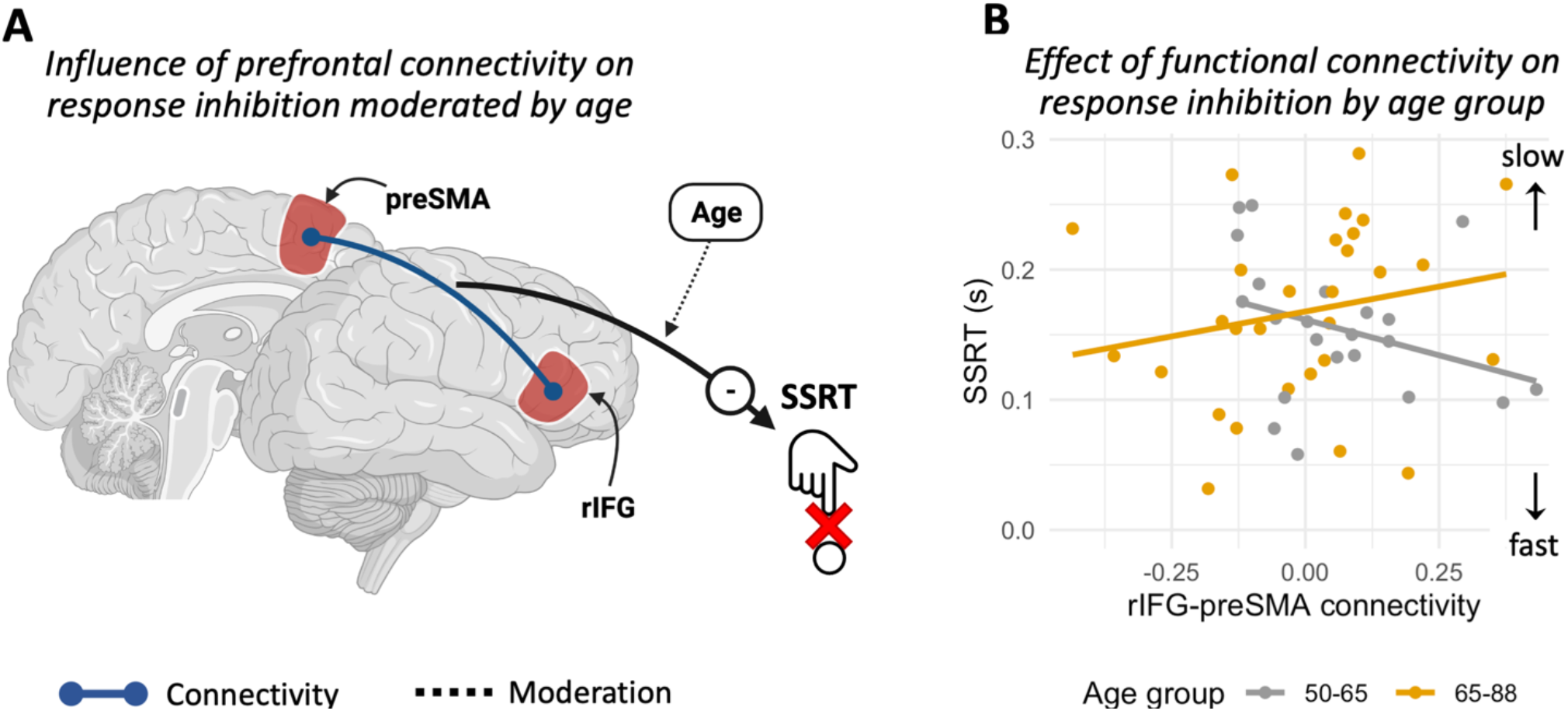
A) Age moderates the influence of connectivity between the preSMA and rIFG on response inhibition (SSRT). B) SSRT estimates as a function of connectivity between preSMA and rIFG. The interaction with age is estimated as a continuous variable (see text) but binarized for visualization purposes only.

### Modulation of Prefrontal connectivity partially mediates influence of locus coeruleus on response inhibition

Next, we tested whether task-related functional connectivity mediated the variance between locus coeruleus signal and SSRT, subject to moderation by age (Figure 4A). The direct association between locus coeruleus and response inhibition was moderated by age (c2: standardized β = 0.817; p = 0.013). The association between connectivity and response inhibition was also conditional on age (b2: standardized β = 0.637; p < 0.0001). A formal test of moderated mediation based on the index term (Hayes, 2015) indicated that the effect of locus coeruleus integrity on response inhibition was partly mediated by functional connectivity in the prefrontal stopping-network, but this relationship changed with age (ab: coefficient of moderated mediation = 0.00019; bootstrapped 95% CI = [0.00003, 0.00053]; p = 0.004; proportion moderated = 0.159). The moderated direct and indirect effects of locus coeruleus integrity on response inhibition are depicted on figure 4B.

**Figure 4.**
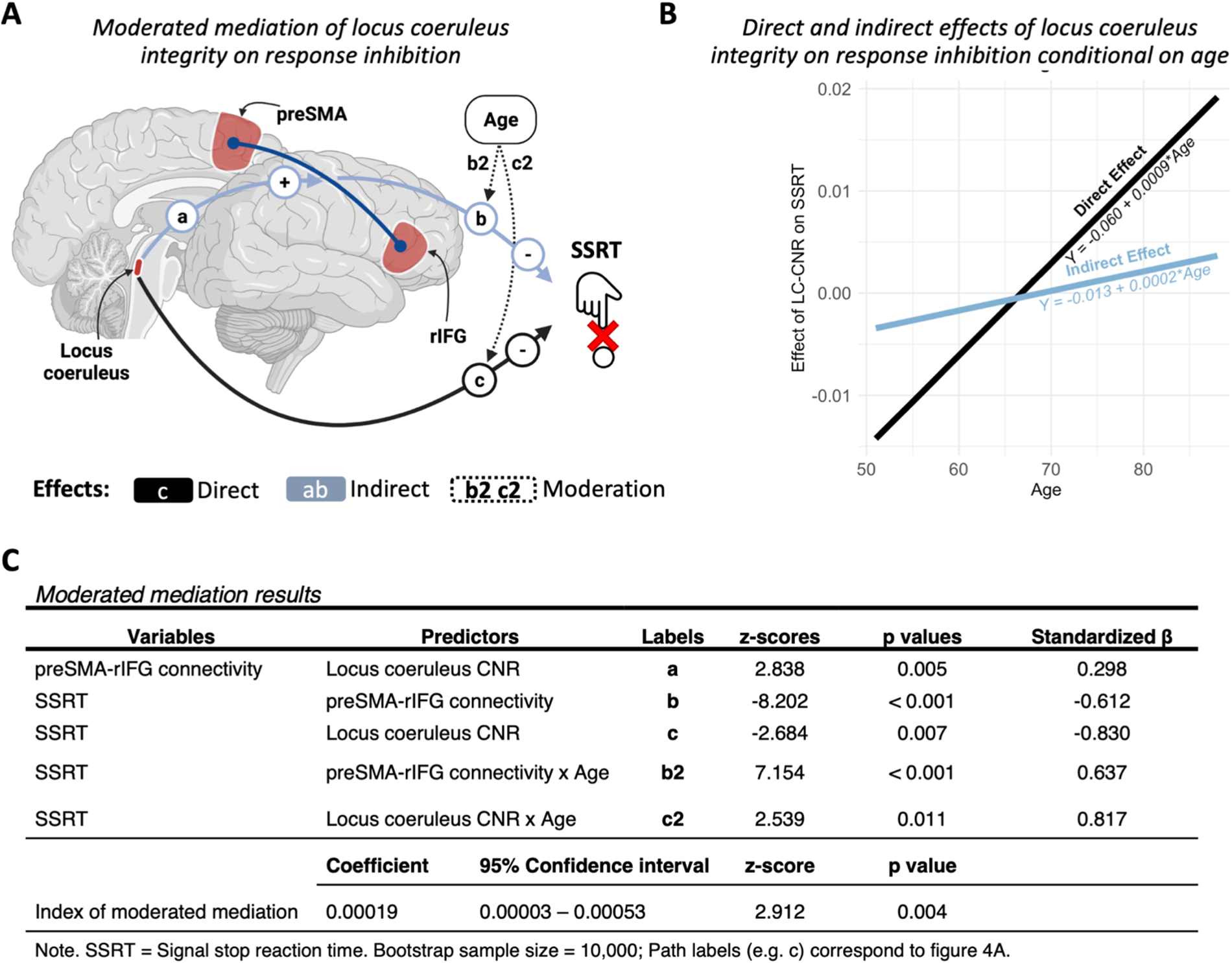
A) Moderated mediation model, with locus coeruleus integrity as a predictor (LC-CNR), functional connectivity within the prefrontal stopping-network as a mediator (Connectivity), and stop signal reaction times as an outcome (SSRT). Age was included as a second-level moderator. B) Direct and indirect (i.e., mediated by functional connectivity) effects of locus coeruleus integrity on response inhibition, conditional on age.

## Discussion

The principal results of this pre-registered study are that (i) individual differences in stopping efficiency are related to the integrity of the locus coeruleus; and (ii) this effect is partially mediated by the facilitation of connectivity within the prefrontal stopping-network. These effects were observed in healthy adults from a population-based cohort, but moderated by age, over and above the main effect of age on locus coeruleus integrity.

This behavioural effect is mainly associated with integrity of the rostral sub-region of the locus-coeruleus, which preferentially targets prefrontal cortex, and is affected by healthy ageing (Loughlin et al., 1986; Mason et al., 1979). Crucially, these effects were observed after correction for a possible main effect of age on the locus coeruleus. Nonetheless, the influence of locus coeruleus variability is moderated by age. Previous reports linked variation of rostral locus coeruleus intensity to cognitive performances in healthy subjects. Preserved integrity of the rostral locus coeruleus in healthy older adults has been associated with better cognitive function (Manaye, McIntire, Mann, and German, 1995; Hammerer et al., 2018; Dahl et al., 2019; Liu et al., 2020). Our results support this previous work and extend the evidence for an influence of the locus coeruleus noradrenaline system on inhibitory control.

The influence of noradrenaline may also be understood in terms of its effect on task-related connectivity between regions (Rae et al., 2016; Holland, Robbins, & Rowe, 2021). For inhibitory control, connectivity between the rIFG and preSMA are of particular relevance: lesion studies, transient interference by magnetic or electrical stimulation, and neuroimaging work provide converging evidence that interactions between these regions are crucial for successful response inhibition (Chambers et al., 2006; Duann et al., 2009; Forstmann et al., 2012; Aron et al., 2014; Rae et al., 2015). We found that the effect of locus coeruleus integrity on response inhibition is partly mediated by changes in functional connectivity between the rIFG and preSMA. These findings reinforce previous reports on the effect of the noradrenaline reuptake inhibitor atomoxetine on response inhibition in Parkinson’s disease. For example, Rae et al (2016) observed that in patients, connectivity between of preSMA on the rIFG was reduced, but restored by atomoxetine. Using dynamic causal modelling, they showed that atomoxetine exerts restored the effective connectivity between preSMA and rIFG, and increased the strength their interacting projections to subthalamic nucleus. Our results similarly associate increased connectivity between these two areas to enhanced response inhibition, and provide evidence that modulation within the stopping-network is related to the integrity of the locus coeruleus, the principal source of noradrenaline.

The influence of both locus coeruleus integrity and prefrontal stopping-network connectivity on inhibitory control changed with age. Our results confirmed such an interaction: higher age-adjusted integrity within the younger and middle age range (50-65 years) of healthy adults was associated with positive modulation of the stopping-network and better inhibitory control. This was not observed over 65 years.

Ageing is also associated with decline of other aspects of motor performance such as psychomotor slowing and reduced fine motor skills (Salthouse, 2000; Seidler et al., 2010). A contributor to age-related performance decline is loss of grey matter volume (Draganski, Lutti, & Kherif, 2013) with evidence from functional neuroimaging studies for adaptive plasticity paralleling structural decline (Tsvetanov et al., 2020). Consequently, older adults may display more widespread brain activation, weaker segregation of local networks and weaker inter-hemispheric connectivity (Rowe et al., 2006; Chan, Park, Savalia, Petersen, & Wig, 2014; Geerligs, Renken, Saliasi, Maurits, & Lorist, 2014; Tsvetanov et al., 2016). Evidence to date suggests that less segregated brain networks contribute to age-related decline in cognitive and sensorimotor performance (Chan, Park, Savalia, Petersen, and Wig, 2014; Geerligs, Renken, Saliasi, Maurits, and Lorist, 2014; King et al., 2017; Bethlehem et al., 2020). Indeed, similarly to our results, analysis of cortical connectivity in the CamCAN dataset showed that stopping efficiency in older adults relied more strongly on connectivity than in their younger counterparts (Tsvetanov et al., 2018), in accord with preclinical studies of the effects of locus coeruleus plasticity and connectivity (Bear & Singer, 1986; Coull, Buchel, Friston, Frith, 1999; Martins & Froemke, 2015). Further longitudinal studies are warranted to investigate age-related changes in the way locus coeruleus integrity and cortical connectivity impact on inhibitory control.

Our pre-registered analysis was designed to probe the association of locus coeruleus integrity with response inhibition and differs in many respects from an earlier analysis of the Cam-CAN cohort looking for associations of coeruleus integrity with diverse cognitive functions (Liu et al., 2020). We focused on adults aged above 50 years, for better estimation of structural integrity of the locus coeruleus using MT weighted MRI. This is because of the greater neuromelanin based contrast with age (Zecca, Youdim, Riederer, Connor, & Crichton, 2004). The locus coeruleus signal was extracted through a probabilistic atlas-based segmentation, which provides unbiased estimation, with superior accuracy and reliability compared to the manual segmentation approaches (Chen et al., 2014; Langley, Huddleston, Liu, & Hu, 2016; Ye et al., 2021). These features are preferable when using 3T MRI images, where the manual segmentation approach may be unreliable due to the relatively low signal-to-noise ratio. Next, the stop-signal reaction time (SSRT) was estimated using a Bayesian parametric model of the stop-signal task which isolates attentional confounds from response inhibition (Matzke, Dolan, Logan, Brown, & Wagenmakers, 2013). These differences between studies might explain the fact that no effect of locus coeruleus signal on the stop-signal task was observed in Liu et al (2020).

There are limitations in the current study that should be considered when interpreting the results and need to be addressed in future studies. First, the focus on ages above 50 years reduced sample size. Power calculations confirmed that our study was well powered for the intended analyses (see pre-registration <https://osf.io/zgj9n/>). However, individual variability in locus coeruleus signal increases with age (Liu et al., 2019; Ye et al, 2020). Second, our results are based on a cross-sectional cohort. Therefore, our conclusions merely speak to the effects of age and its correlates, as assessed across individuals, but provide no insight on the dynamic process of individual ageing. A larger sample size including more subjects of advanced age or a longitudinal cohort might be needed in future studies to confirm the impact of ageing on the relationship between locus coeruleus and inhibitory control. Third, participants’ cognitive functions were screened with the ACE-R test which may lack the sensitivity to detect latent Alzheimer’s disease pathology in the older group. Notably, the regional vulnerability of locus coeruleus neurons varies between disorders and the rostral region, which we show being related to response inhibition, is especially vulnerable to Alzheimer’s disease and latent pathology with healthy ageing (German et al., 1992; Mason & Fibiger, 1979). Fourth, the 3T MRI images used in the present study afford a lower signal-to-noise ratio compared to 7T scans. However, the resulting reduced sensitivity would be expected to increase type II but not type I error. Finally, the present work focused on the modulatory role of the locus coeruleus noradrenergic system on action cancellation. We acknowledge that other neuromodulators, such as serotonin, can play a crucial role in regulating forms of inhibitory control other than action cancellation (Eagle, Bari, and Robbins 2008; Cools, Roberts, and Robbins, 2008; Ye et al. 2014; Hughes, Rittman, Regenthal, Robbins, & Rowe 2015).

In conclusion, we show that the ability to inhibit responses relies on both the locus coeruleus and its facilitation of connectivity within the prefrontal cortex. The locus coeruleus integrity has different implications for inhibitory control at different ages. These findings contribute to the broader understanding of the importance of noradrenergic systems for executive functions in normal populations, with implications for impulsive clinical disorders.

## Acknowledgments

This work was funded by a Medical Research Council intramural grant (SUAG/051 R101400) and a Wellcome Trust grant (220258) awarded to J.B.R.; K.A.T. was supported by the Guarantors of Brain (G101149). The authors declare no competing financial interests and no conflict of interest.

